# Identification of stripe rust adult plant resistance genes in the hard winter wheat cultivar Baker’s Ann

**DOI:** 10.1101/2025.09.04.674321

**Authors:** Rajat Sharma, Meinan Wang, Xianming Chen, Brett F. Carver, Mary Guttieri, Paul St. Amand, Amy Bernardo, Guihua Bai, Anju Maan Ara, Katherine W. Jordan, Meriem Aoun

**Affiliations:** Department of Entomology and Plant Pathology, Oklahoma State University, Stillwater, OK, USA; Department of Plant Pathology, Washington State University, Pullman, WA, USA; USDA-ARS Wheat Health, Genetics, and Quality Research Unit, Pullman, WA, USA; Department of Plant and Soil Sciences, Oklahoma State University, Stillwater, OK, USA; USDA-ARS Hard Winter Wheat Genetics Research Unit, Manhattan, KS, USA; Department of Agronomy, Kansas State University, Manhattan, KS, USA

## Abstract

Stripe rust, caused by *Puccinia striiformis* f. sp. *tritici* (*Pst*), is among the most destructive wheat diseases. Identifying resistance genes is crucial for the development of resistant cultivars. “Baker’s Ann”, a hard winter wheat cultivar developed by Oklahoma State University, has shown durable adult plant resistance to stripe rust. To dissect the genetic basis underlying stripe rust resistance in Baker’s Ann, 125 doubled haploid lines, derived from the cross OK12D22004-016 × Baker’s Ann, were evaluated at the adult plant stage in the greenhouse and in field environments in Oklahoma, Kansas, and Washington. This population was genotyped using genotyping-by-sequencing, which produced 7,268 single-nucleotide polymorphisms for genetic mapping. Quantitative trait loci (QTL) analysis identified six loci, four from Baker’s Ann on chromosomes 2DL, 4BS, 4BL, and 7BL, and two from OK12D22004-016 on chromosomes 2AS and 2AL. Although OK12D22004-016 is susceptible in the US Great Plains, it was found to carry *QYr.osu-2AS*, which was linked to *Yr17* on the 2N^v^S translocation and explained up to 30% of the phenotypic variation, but was effective in a single location in Washington. Two major QTL were identified in Baker’s Ann, *QYr.osu-2DL* on chromosome 2DL that explained up to 57% of the phenotypic variation and mapped close to *Yr54*, and *QYr.osu-4BL* on chromosome 4BL that explained up to 15% of the phenotypic variation and mapped close to *Yr62*. Resistance in Baker’s Ann resulted from additive effects of the four QTL. Two kompetitive allele-specific PCR markers were developed for *QYr.osu-2DL* to facilitate marker-assisted selection for stripe rust resistance.

## Core Ideas

- The hard winter wheat cultivar Baker’s Ann is highly resistant to stipe rust at the adult plant stage.
- Four quantitative trait loci (QTL) were mapped in Baker’s Ann on chromosomes 2DL, 4BS, 4BL, and 7BL.
- *QYr.osu-2DL* and *QYr.osu-4BL* were major QTL in Baker’s Ann and mapped close to *Yr54* and *Yr62*, respectively.
- Kompetitive allele specific PCR markers developed for *QYr.osu-2DL* should be useful in marker-assisted selection.

## Introduction

Wheat (*Triticum aestivum* L.) is a staple crop that feeds around 36% of the global population (Talha et al., 2016) and contributes about 40% of total caloric intake (Li et al., 2019). Among various market classes, winter wheat comprises about 70% of total United States (US) production, with hard winter wheat (HWW) predominantly grown in the US Great Plains (USDA Economic Research Service, 2024). Stripe rust (also known as yellow rust), caused by the biotrophic fungus *Puccinia striiformis* f. sp. *tritici* (*Pst*), is one of the most damaging diseases of wheat worldwide (Wellings, 2011). Approximately 88% of the global wheat area is vulnerable to stripe rust, with annual losses exceeding USD 1 billion (Beddow et al., 2015). Stripe rust disease can cause significant yield losses, reaching up to 100%, depending on the cultivar, timing of infection, and environmental conditions (Chen, 2005). Historically, *Pst* isolates thrived in cool and humid climates. Isolates emerging after 2000 have become more aggressive and better adapted to the warmer conditions prevalent in the Great Plains (Chen et al., 2010; Markell & Milus, 2008; Milus et al., 2009). Although foliar fungicides can manage stripe rust, their use increases production costs and raises environmental concerns. Thus, growing resistant wheat cultivars remains the most effective, economical, and environmentally sustainable management strategy (Chen, 2005).

Stripe rust resistance (*Yr*) genes are classified, based on their effectiveness at different plant growth stages, into all-stage resistance (ASR) and adult plant resistance (APR). ASR, also known as seedling resistance, is usually race-specific, qualitatively inherited, effective throughout all growth stages, and typically provides complete resistance. However, ASR genes exert strong selection pressure on the pathogen, making them vulnerable to becoming ineffective with the evolution of new virulent races. Therefore, deploying ASR alone is not recommended to achieve durable resistance. Conversely, APR is usually non-race specific, quantitatively inherited, provides partial but durable resistance, though race-specific APR genes have also been identified (Milus et al., 2015). Cultivars with APR alone remain susceptible at the seedling stage but gradually develop resistance as they mature (Chen, 2013; Ellis et al., 2014; Lagudah, 2011; Mundt, 2014). Nevertheless, a single APR gene typically offers insufficient protection under severe epidemics (Chen, 2014; Risk et al., 2012; Singh et al., 2015). However, near or complete resistance can be achieved by pyramiding multiple APR genes (Sørensen et al., 2014). Hence, combining multiple effective ASR and APR genes or multiple APR genes in a wheat cultivar is recommended to achieve durable stripe rust resistance (Sharma et al., 2025).

To date, 87 officially designated *Yr* genes, 77 temporary named genes, and over 900 quantitative trait loci (QTL) have been identified in wheat and its wild relatives (McIntosh et al., 2020; Sharma et al., 2024, 2025). Of the 87 *Yr* genes, 27 genes confer APR, including *Yr11*, *Yr12*, *Yr13*, *Yr14*, *Yr16*, *Yr18*, *Yr29*, *Yr30*, *Yr36*, *Yr39*, *Yr46*, *Yr48*, *Yr49*, *Yr52*, *Yr54*, *Yr55*, *Yr58*, *Yr59*, *Yr62*, *Yr68*, *Yr71*, *Yr75*, *Yr77*, *Yr78*, *Yr79*, *Yr80*, and *Yr86*, and the other 60 genes confer ASR. Eleven *Yr* genes have been cloned, of which eight are ASR genes, namely *Yr5* (*YrSP)*, *Yr7*, *Yr10* (*YrNAM*), *Yr15*, *Yr27*, *YrAS2388*, *YrU1*, and *Yr87* (Athiyannan et al., 2022; Klymiuk et al., 2018; Liu et al., 2014; Marchal et al., 2018; Ni et al., 2023; Sharma et al., 2024; Wang et al., 2020; Zhang et al., 2019), and only three are APR genes (*Yr18*, *Yr36*, and *Yr46*). ASR genes typically encode nucleotide-binding site leucine-rich repeat (NBS-LRR) proteins that recognize *Pst* effectors to trigger the resistance response. Proteins encoded by APR genes are unrelated to NBS-LRR proteins. For instance, *Yr18* encodes a putative ATP-binding cassette (ABC) transporter, facilitating sinapyl alcohol export across cell membrane for enhanced cell wall lignification (Krattinger et al., 2009; Zhang et al., 2024), *Yr36* encodes a protein kinase (WKS1) (Fu et al., 2009), and *Yr46* encodes a hexose transporter (Moore et al., 2015). Despite numerous identified *Yr* genes in wheat, most ASR genes and some APR genes are no longer effective (Hovmøller et al., 2011). Sharma et al. (2025) reported that stripe rust resistance in HWW from the US Great Plains is primarily conferred by APR however, the underlying *Yr* genes are largely unknown. Thus, identifying and characterizing *Yr* genes are critical to diversify resistance sources in wheat cultivars to achieve durable resistance.

“Baker’s Ann” (pedigree: TX00D1390/OK03522; experimental line: OK13621) is a hard red winter wheat cultivar released in 2018 by the Oklahoma State University (OSU) wheat breeding program. Owing to its durable stripe rust resistance, agronomic superiority, and exceptional dough strength, Baker’s Ann has been extensively used in the OSU wheat breeding program. Although susceptible at the seedling stage, Baker’s Ann has consistently demonstrated stable and high-level stripe rust resistance at the adult plant stage since 2014. However, the genetic basis of this APR remains unclear. The objectives of this study were to (1) identify and map QTL associated with stripe rust APR in Baker’s Ann using single-nucleotide polymorphisms (SNPs) generated through genotyping-by-sequencing (GBS), and (2) develop kompetitive allele-specific PCR (KASP) markers from tightly linked SNPs to the resistance QTL for use in marker-assisted selection (MAS).

## Materials and Methods

### Plant materials

A doubled haploid (DH) population consisting of 125 lines derived from the bi-parental cross OK12D22004-016 × Baker’s Ann was used in this study. OK12D22004-016 (pedigree: Everest/OK08328//OK09634) is an OSU experimental line that is susceptible to stripe rust at the seedling stage and moderately susceptible at the adult plant stage in Oklahoma fields. Baker’s Ann is susceptible at the seedling stage to US *Pst* races but exhibits a strong APR in multiple environments in the US since 2014. Stripe rust-susceptible checks “Jagalene”, “Pete”, and “PS 279” were included in this study to monitor disease pressure, with at least one check planted in each test environment.

### Stripe rust phenotyping at the adult plant stage

A total of 125 DH lines of the cross OK12D22004-016 × Baker’s Ann, the two parents, and two susceptible checks (Pete and Jagalene) were evaluated for reaction to race PSTv-37 at the flag-leaf stage under controlled greenhouse conditions. PSTv-37 is the most predominant race in the US with virulence to *Yr6*, *Yr7*, *Yr8*, *Yr9*, *Yr17*, *Yr27*, *Yr43*, *Yr44*, *Yr85* (= *YrTr1*), and *YrExp2* among the 18 genes in the *Yr* single-gene lines used to differentiate *Pst* races (Feng et al., 2023; Wan & Chen, 2014). Six seedlings per line were vernalized at 4°C under a 12-h photoperiod for 6 – 8 weeks. After vernalization, seedlings were transplanted into cone-tainers (SC10U UV-stabilized cones, 3.8 cm diameter × 20.9 cm depth, 164 mL; Stuewe & Sons, Tangent, OR) with two plants per cone-tainer. Each line had six plants allocated into three independent replications, with one cone-tainer representing each replicate. Susceptible checks and parents were planted after every 20 DH lines. The plants were grown at 22°C/18°C (day/night) with a 16-h photoperiod. At the flag leaf stage, plants were uniformly inoculated with urediniospores suspended in Soltrol 170 mineral oil (Chevron Phillips Chemical Company, The Woodlands, TX) at a concentration of 10 mg mL^−1^. Following inoculation, plants were air-dried and incubated for 24 hours in a dark humidity chamber (10°C, 100% relative humidity), then transferred to a rust-free growth chamber maintained at 20°C/4°C (day/night) with a 16-h photoperiod. Infection type (IT) was recorded 18–21 days post-inoculation on a 0–9 scale, with 0 – 3 indicating resistance, 4 – 6 moderate resistance, and 7 – 9 susceptibility (Line & Qayoum, 1992; Wan et al., 2017). The mean IT across three replications per line was used for further analysis.

The field experiments were conducted for stripe rust responses in 2024 across four locations: Chickasha in Oklahoma, Rossville in Kansas, Pullman and Mount Vernon in Washington. These four locations are spread over wide geographically region in US and have different weather conditions and *Pst* race compositions. The DH lines were planted in Fall 2023 with two replications at Chickasha and one replication with repeated checks at each of the other locations. The lines were planted in 1.2 m rows.

The nurseries at Chickasha and Rossville were artificially inoculated with local races specific for each location to ensure proper disease development. The Pullman and Mount Vernon fields were naturally infected. Standard management practices were adopted in each location. At Chickasha, susceptible checks Pete and Jagalene, along with parents, were planted after every 50 DH lines and Pete was used as spreader and border rows. Stripe rust responses were recorded on flag leaves using IT on a 0–9 scale and disease severity (DS) based on the percentage of infected flag leaf area (Peterson et al., 1948). The disease ratings (IT and DS) were collected twice at Feekes stage 10.5 (Large, 1954) on April 30 (CH 1) and May 4 (CH 2). In Pullman and Mount Vernon, Jagalene and PS 279 were used as susceptible checks, and PS 279 was also planted as border rows. In Pullman, disease ratings were taken at Feekes stage 10.5 on June 12 (PL). In Mount Vernon, data were recorded two times, first at Feekes stage 4 (early jointing stage) on April 10 (MV 1) and second at Feekes stage 10.54 (kernel watery ripe) on June 6 (MV 2). In Rossville, Jagalene was used as susceptible check and planted after every 80 rows. In Rossville, data for IT and DS were taken at Feekes stage 10.5 on May 17 (RS). Best linear unbiased estimates (BLUE) for IT and DS were also calculated across environments using R package “lme4” (Bates et al., 2015; Vazquez et al., 2010) as described by Aoun et al. (2022), where line was considered as a fixed effect and environment was considered as a random effect. Each IT and DS rating from individual locations, along with BLUE and greenhouse ratings, was treated as a distinct trait. The abbreviations for the different traits are defined in Table 1 and used hereafter. Analysis of variance (ANOVA) for IT and DS data was conducted using the “aov” function in R to assess genotype and environment effects. Pearson’s correlation for stripe rust responses across environments was performed using ‘metan’ package in R (Olivoto & Lúcio, 2020).

**Table 1.** Stripe rust responses in different environments for the doubled haploid (DH) population of the cross OK12D22004-016 × Baker’s Ann and its parents.

| Trait/environment <sup>a</sup> | Designation | Parents |  | DH population |  |
| --- | --- | --- | --- | --- | --- |
|  |  | OK12D22004-016 | Baker's Ann | Range | Mean ± SE <sup>b</sup> |
| IT–Greenhouse (4–20 °C) | GH (IT) | 8 | 2 | 1 – 9 | 5.49 ± 0.23 |
| IT–Chickasha, OK (April 30, 2024) | CH 1 (IT) | 6 | 1 | 1 – 9 | 4.02 ± 0.23 |
| IT–Chickasha, OK (May 4, 2024) | CH 2 (IT) | 6 | 2 | 1 – 9 | 4.09 ± 0.22 |
| IT–Rossville, KS (May 17, 2024) | RS (IT) | 7 | 2 | 0 – 9 | 4.67 ± 0.24 |
| IT–Pullman, WA (June 12, 2024) | PL (IT) | 9 | 2 | 2 – 9 | 5.54 ± 0.24 |
| IT–Mount Vernon (April 10, 2024) | MV 1 (IT) | 5 | 4 | 2 – 9 | 4.92 ± 0.25 |
| IT–Mount Vernon (June 6, 2024) | MV 2 (IT) | 3 | 2 | 2 – 9 | 4.95 ± 0.26 |
| DS–Chickasha, OK (April 30, 2024) | CH 1 (DS) | 33 | 3 | 1 – 80 | 19.90 ± 1.99 |
| DS–Chickasha, OK (May 4, 2024) | CH 2 (DS) | 48 | 6 | 1 – 83 | 23.50 ± 2.23 |
| DS–Rossville, KS (May 17, 2024) | RS (DS) | 48 | 3 | 0 – 95 | 29.90 ± 2.47 |
| DS–Pullman, WA (June 12, 2024) | PL (DS) | 98 | 10 | 2 – 100 | 43.10 ± 2.85 |
| DS–Mount Vernon (April 10, 2024) | MV 1 (DS) | 50 | 35 | 10 – 70 | 36.88 ± 1.856 |
| DS–Mount Vernon (June 6, 2024) | MV 2 (DS) | 18 | 10 | 5 – 100 | 48.32 ± 3.23 |
| IT–Multi-environment BLUE | BLUE (IT) | 6 | 2 | 1 – 9 | 5.06 ± 0.21 |
| DS–Multi-environment BLUE | BLUE (DS) | 52 | 13 | 4.2 – 86.6 | 36.34 ± 2.25 |

<sup>a</sup> IT = Infection type; DS = Disease severity; BLUE = Best linear unbiased estimates; Traits at MV1 (Mount Vernon, April 10) were evaluated at early jointing stage, while others were evaluated at the flowering stage.
<sup>b</sup> SE, Standard error of the mean.

### Genotyping of the DH population

Leaf tissues of the DH lines and the parents were collected at the two-leaf stage in 96-well plates and ground into a fine powder using one 4.5 mm steel bead per well and shaking at 1,750 cycles/min for 3 min in a 2010 Geno/Grinder (Spex SamplePrep LLC, Metuchen, NJ). DNA was extracted using a modified cetyltrimethylammonium bromide (CTAB) approach developed by the USDA Central Small Grain Genotyping Lab, Manhattan, KS (Serba et al., 2019). The DH population and the parents were genotyped using GBS (Elshire et al., 2011; Poland et al., 2012) at the USDA-ARS Genotyping Lab in Manhattan, KS. The raw sequencing data can be accessed from the NCBI Short-Read Archive BioProject PRJNA1312051 (ncbi.nlm.nih.gov/sra/PRJNA1312051). The SNP calling was performed in TASSEL software v.5 (Bradbury et al., 2007), and marker physical positions were assigned according to the Chinese Spring reference genome IWGSC RefSeq v2.1 (Zhu et al., 2021). GBS-SNP markers used for downstream analysis are available in Table S1. To ensure data quality, SNP with missing data ≥ 40%, minor allele frequency ≤ 5%, or heterozygosity ≥ 15% were removed. The remaining 13,087 high-quality SNPs were sorted according to parents, and all monomorphic markers or markers for which either parent was heterozygous or had missing data were removed. This resulted in 7,268 polymorphic SNP markers, which were used for subsequent analysis (Table S2). After identifying chromosomes containing resistance QTL, simple sequence repeats (SSR), and KASP markers linked to known *Yr* genes or QTL in proximity to the identified major QTL in our study were tested on parental lines. Polymorphic markers were then used to genotype the DH population, and the resulting data were merged with the original GBS SNP markers (Table S2) to refine the linkage map. Primer details for KASP and SSR markers, along with their PCR conditions are provided in Table S3.

### Linkage mapping and QTL analysis

The high-quality SNP markers were further filtered using “BIN” functionality in QTL IciMapping v4.2 (Meng et al., 2015) to remove the redundant markers (with same genetic positions). The “BIN” functionality was also used to remove markers showing segregation distortion based on Chi-squared test that did not fit the 1:1 segregation ratio at a *P* value ≤ 0.01. The resulting data was examined for the presence of switch alleles, genotyping errors, excessive crossover count and double crossover using the R package “R/qtl” (Broman, 2010; Broman et al., 2003). Then, linkage map was constructed using “MAP” functionality in QTL IciMapping v4.2. The markers were grouped into linkage groups using minimum logarithm of odds (LOD) score of 6.0 and, generated linkage groups were assigned to corresponding chromosomes based on marker physical positions on the reference genome IWGSC RefSeq v2.1 (Zhu et al., 2021). Within each linkage group, markers were ordered with the 2-OptMap algorithm and refined with the ripple function with a window size of 5. Recombination frequencies were converted to genetic distance in centiMorgan (cM) using the Kosambi mapping function (Kosambi, 1943).

The linkage map and phenotypic data across environments were used to identify the resistance QTL. QTL analysis was conducted using “BIP” functionality and inclusive composite interval mapping for additive QTL (ICIM-ADD) method in QTL IciMapping v4.2. The parameters were set at walk speed of 1 cM and PIN (probability of marker included in model via stepwise regression) of 0.001. The LOD threshold to determine the QTL was calculated using 1000 permutations at *P* = 0.05. QTL detected with at least two traits were reported.

### Marker development and validation

Genomic DNA was isolated from leaf tissue of the two parents and the 125 DH lines at the two-leaf stage using a modified CTAB method adapted for a 96-well plate (Liu et al., 2006; Riede & Anderson, 1996). DNA concentration and purity were assessed with a NanoDrop ND-1000 spectrophotometer (Thermo Scientific, Wilmington, DE, USA). DNA stocks were diluted to a working concentration of 30 ng µL^-1^. After mapping the stripe rust resistance QTL, the closest SNPs to major QTL were selected and converted into KASP markers. As high sequence similarity among homoeologous and paralogous regions in hexaploid wheat can occur, locus-specific common primers were designed using the anchoring strategy recommended by LGC Biosearch Technologies. (https://biosearchassets.blob.core.windows.net/assetsv6/guide_kasp-assay-design-anchoring.pdf). A 150 bp DNA sequence flanking each SNP was blasted to identify an anchor point (unique base) at the 3’ end of the common primer to ensure locus specificity. HEX tail 5′-GAAGGTCGGAGTCAACGGATT-3′ and FAM tail 5′-GAAGGTGACCAAGTTCATGCT3′ were added to 5’ end of allele-specific primers for resistant and susceptible alleles, respectively. Primers were synthesized by Integrated DNA Technologies (Coralville, IA). KASP reactions were performed using the modified LGC Biosearch Technologies thermal-cycling protocol (Makhoul et al., 2020). Each 100 µL primer mix contained 12 µL of each allele-specific primer (100 µM), 30 µL of the common primer (100 µM), and 46 µL of ddH_2_O. Each PCR reaction contained 5 µL of 2× KASP master mix (KBS-1050-102; LGC Biosearch Technologies Hoddesdon, United Kingdom), 0.14 µL primer mix, 1.86 µL of ddH_2_O and 3 µL of DNA (30 ng/µL). The PCR was conducted on a Bio-Rad CFX-96 Opus real-time PCR system (Bio-Rad, Hercules, CA) with the following profile: hot start at 94 °C for 15 min, followed by 10 touchdown cycles at 94 °C for 20 s and 65 °C for 60 s with a decrease of 0.8 °C per cycle, followed by 30 cycles of 94 °C for 20 s and 57 °C for 60 s with final step to read the plate at 37 °C. Endpoint fluorescence and genotype clustering were performed in Bio-Rad CFX Maestro Software 2.3. One to three additional recycling steps were performed if genotype clusters were not clearly differentiated after initial PCR. The recycling step comprised of three additional PCR cycles at 94 °C for 20 s and 57 °C for 60 s. Each marker assay was validated first on the parents and subsequently used to genotype the 125 DH lines.

### Exome Capture

Baker’s Ann, alongside other 82 HWW accessions, was previously genotyped using exome capture generated as part of a project described in Jordan et al. (2022) (NCBI Short-Read Archive BioProjects PRJNA381058 and PRJNA732645) by USDA-ARS Hard Winter Wheat Genetics Research Unit in Manhattan, KansasWheat exome capture was performed as previously described in Krasileva et al (2017). The genomic libraries were sequenced on Illumina NovaSeq instrumentation using 2 × 150 bp read runs as described by Jordon et al. (2022). Raw fastq files were filtered for average read quality ≥20 using bbduk tools v39.01 (Bushnell, 2014). The QC filtered reads were aligned to the Chinese Spring wheat reference genome RefSeq v.2.1 (Zhu et al., 2021) using HISAT2 aligner v2.2.1 (Kim et al., 2019), retaining only uniquely mapped reads. The resulting alignments were processed using samtools v1.17 (Li et al., 2009) and then analyzed using the best practices workflow in Genome Analysis Toolkit (GATK) v4.4.0 (McKenna et al., 2010), which produced a g.vcf file for each sample using GATK’s Haplotype Caller algorithm. A raw single .vcf file was generated using the CombineGVCFs and GenotypeGVCFs tools within GATK v4.4.0. The raw variant file was filtered using Perl to retain biallelic variants with data present in > 80% of HWW accessions. The exome capture SNP data were used to assess the relationship between a major QTL identified in Baker’s Ann and another stripe rust resistance QTL previously identified in the HWW cultivar “Overland” (Mustahsan et al., 2023). Therefore, exome capture SNP data were further filtered to retain SNPs that were located within the physical positions of the major resistance QTL in Baker’s Ann. Kinship, based on identity-by-state, and UMPGA (Unweighted Pair-Cluster Method using Arithmetic Averages) tree in the selected region were performed using Tassel v5.2.96 (Bradbury et al., 2007).

## Results

### Phenotypic evaluation

Baker’s Ann displayed high resistance at the adult plant stage, with IT ranging from 0 – 2 and DS between 3 – 13% across environments. However, at the early jointing stage at MV 1, it exhibited moderate resistance (IT 4, DS 35%) (Table 1; Figure 1 and 2). OK12D22004-016 varied from moderately resistant to susceptible (IT 5–9, DS 33–98%) across environments, though it was resistant at MV 2 (IT 3, DS 18%). Mean IT values for the DH lines across environments ranged from 4.0 to 5.5, whereas mean DS values ranged from 19.9 to 48.3% (Table 1). The distributions for IT and DS at the adult plant stage were in most environments continuous (Figure 1 and 2; Table S4), suggesting that resistance was likely governed by multiple QTL. As OK12D22004-016 was moderately resistant in some environments, transgressive segregants were also observed. ANOVA for IT and DS data showed significant effects for the DH lines and environments (Table S5). Significant positive correlations (*P* < 0.001) were found for IT and DS data across all environments (Figure S1).

**Figure 1.**
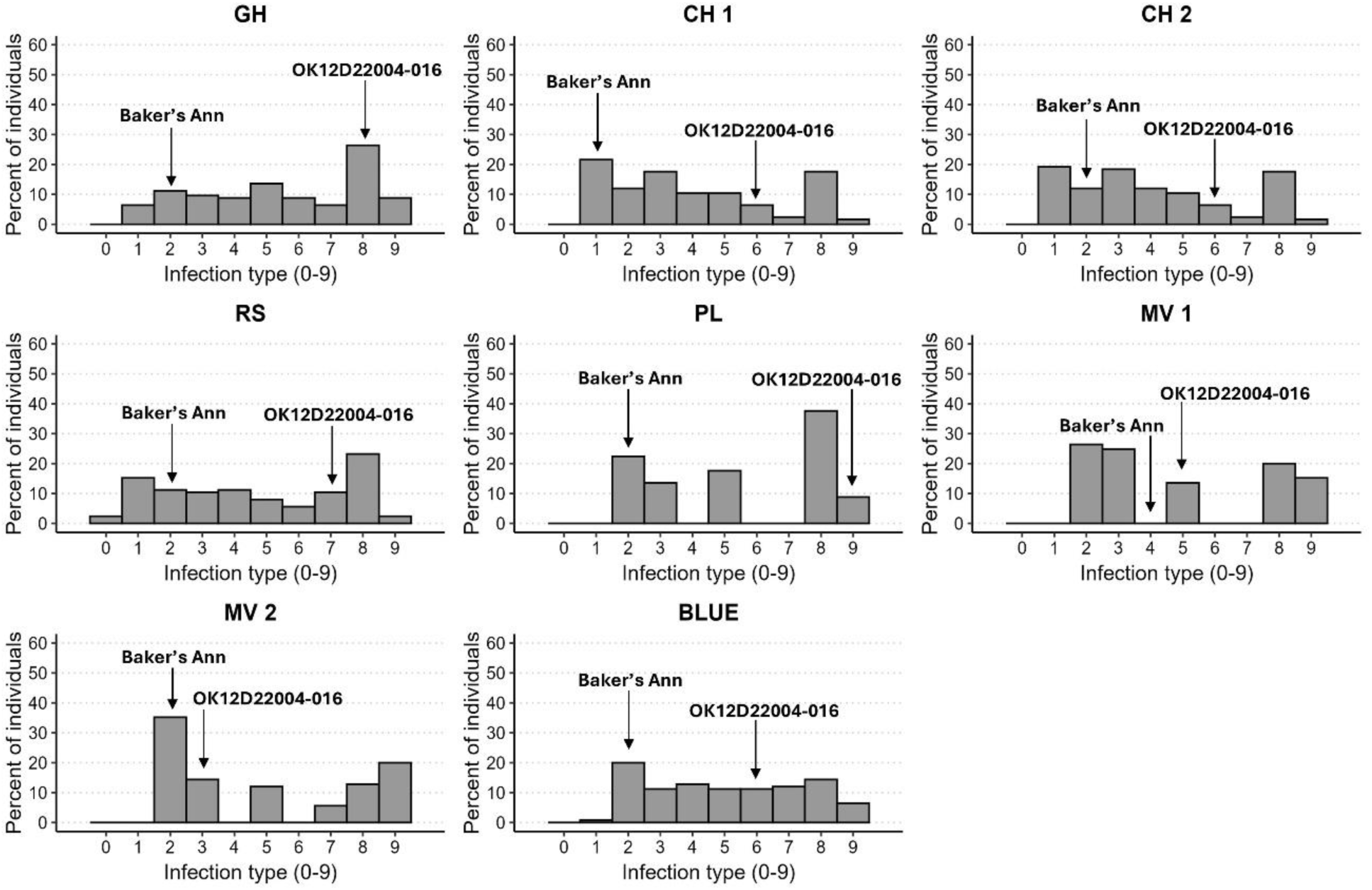
Distribution of stripe rust infection type at the adult plant stage of 125 doubled haploid lines from the cross OK12D22004-016 × Baker’s Ann across multiple environments. GH = Greenhouse (4– 20°C); CH 1 = Chickasha, OK (evaluated on April 30), CH 2 = Chickasha, OK (evaluated on May 4); RS = Rossville, KS (evaluated on May 17); PL = Pullman, WA (evaluated on June 12); MV 1 = Mount Vernon (evaluated on April 10); MV 2 = Mount Vernon (evaluated on June 6); BLUE = Multi-environment best linear unbiased estimates.

**Figure 2.**
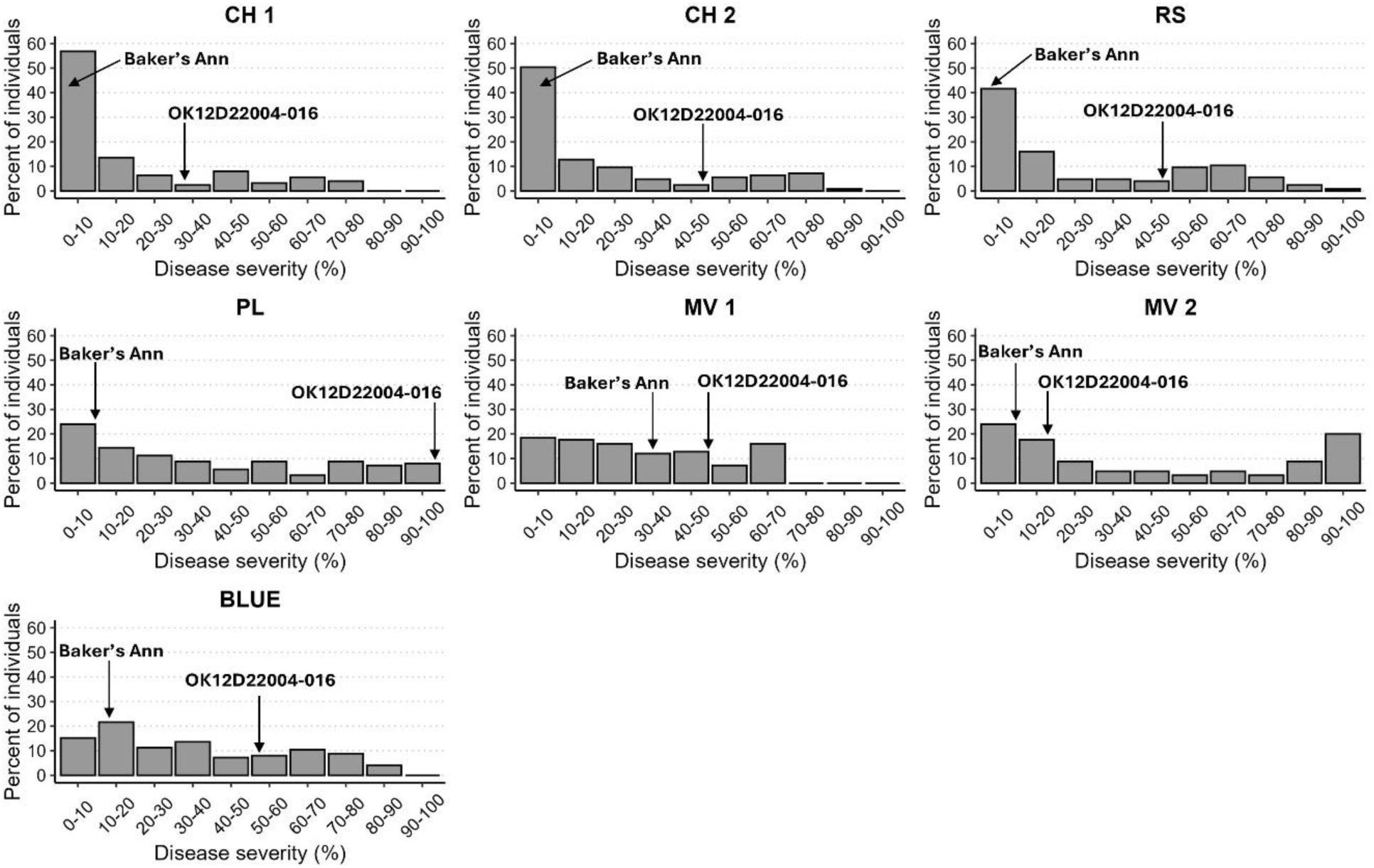
Distribution of stripe rust disease severity at the adult plant stage of 125 doubled haploid lines from the cross OK12D22004-016 × Baker’s Ann across multiple environments. CH 1 = Chickasha, OK (evaluated on April 30), CH 2 = Chickasha, OK (evaluated on May 4); RS = Rossville, KS (evaluated on May 17); PL = Pullman, WA (evaluated on June 12); MV 1 = Mount Vernon (evaluated on April 10); MV 2 = Mount Vernon (evaluated on June 6); BLUE = Multi-environment best linear unbiased estimates.

### Genetic linkage map

After filtering redundant markers and those deviating from the expected 1:1 segregation ratio at *P* > 0.01, 656 SNP markers of the 7,268 polymorphic SNP markers were retained and used for QTL mapping. No markers were excluded for switched alleles or double crossovers, nor were DH lines discarded due to excessive crossover counts. After identifying major QTL, five SSR/KASP markers, *Lr37-Yr17-Sr38_GBG-KASP* (Liu et al., 2020), *Xgwm301* (Basnet et al., 2014), *KASP_2D:638369560* (Mustahsan et al., 2023), *KASP_2D:639534738* (Mustahsan et al., 2023), and *Xgwm251* (Lu et al., 2014), which were previously linked to known *Yr* genes/QTL within the genomic regions of identified QTL in this study, and showed polymorphism between the parents of the DH population were used to genotype the DH lines and integrated in the final linkage map.

The final linkage map consisted of 661 markers distributed across 27 linkage groups, representing all 21 wheat chromosomes (Table S6 and S7). These markers covered a total length of 3,023.4 cM, with an average interval of 4.57 cM between markers. Chromosomes 1B, 2A, 6D, and 7B were each split into two linkage groups, chromosome 2B into three groups, whereas other chromosomes had single groups. The A, B, and D genomes included 286, 233, and 142 markers, respectively, spanning lengths of 1,176.2 cM, 1,000.5 cM, and 846.6 cM. The number of markers per linkage group ranged from six markers in each of the linkage groups 6D-2 (18.7 cM) and 2B-3 (30.2 cM), to 61 markers in the linkage group 5A (231.6 cM).

### QTL analysis

Six QTL were identified at a threshold LOD score of 3.3 (determined based on 1000 permutations), four of which were from Baker’s Ann (*QYr.osu-2DL*, *QYr.osu-4BS*, *QYr.osu-4BL*, and *QYr.osu-7BL*), and two from OK12D22004-016 (*QYr.osu-2AS* and *QYr.osu-2AL)* (Table 2; Figure 3). *QYr.osu-2DL*, *QYr.osu-4BS*, *QYr.osu-4BL* from Baker’s Ann were identified in multiple environments, whereas the remaining QTL from Baker’s Ann and OK12D22004-016 were identified in single locations. Among Baker’s Ann QTL, *QYr.osu-2DL* was a major QTL mapped on the long arm of chromosome 2D at 636.8–651.2 Mb based on IWGSC RefSeq v2.1 and explained 29.7 to 57.1% of the phenotypic variation. LOD scores for *QYr.osu-2DL* ranged from 20.7 for PL (DS) to 38.2 for BLUE (DS). This QTL was detected across all environments and was mapped to a 21.5 cM region flanked by markers *S2D_636865313* (125.5 cM) and *S2D_651288630* (147.1 cM) with *S2D_649277571* (145.3 cM) and *S2D_651288630* (147.1 cM) being the closet markers to this QTL. The second major QTL from Baker’s Ann, *QYr.osu-4BL*, was mapped on chromosome 4BL at the physical position of 544.5 to 595.3 Mb with the closely linked markers being *S4B_544542616* (51.3 cM) and *S4B_595356335* (54.0 cM). *QYr.osu-4BL* was detected in all environments except in MV 2 (IT and DS) and RS (DS). The LOD scores for this QTL ranged between 5.5 to 10 and explained up to 14.6% of the phenotypic variation. Another minor QTL, *QYr.osu-4BS*, was identified in Baker’s Ann on the short arm of chromosome 4B. It was flanked by markers *S4B_4757668* and *S4B_7266951* (0 – 6.5 cM) andwas significant in RS (IT), MV 2 (IT), CH 1 (DS), CH 2 (DS), MV 2 (DS), and BLUE (DS) where it is explained 2.7 to 7.2% of the phenotypic variation. Based on physical positions of the flanking markers, *QYr.osu-4BS* was within 4.7 to 7.3 Mb. Baker’s Ann was also found to carry a second minor QTL, *QYr.osu-7BL*, located at 718.5 to 751.9 Mb on the long arm of 7B and flanked by the SNP markers *S7B_718501007* and *S7B_751946717* (105.9 – 128.0 cM). *QYr.osu-7BL*was detected for IT and DS in MV 1, MV2, and BLUE, and explained 3.9 to 7.4% of the phenotypic variation.

**Figure 3.**
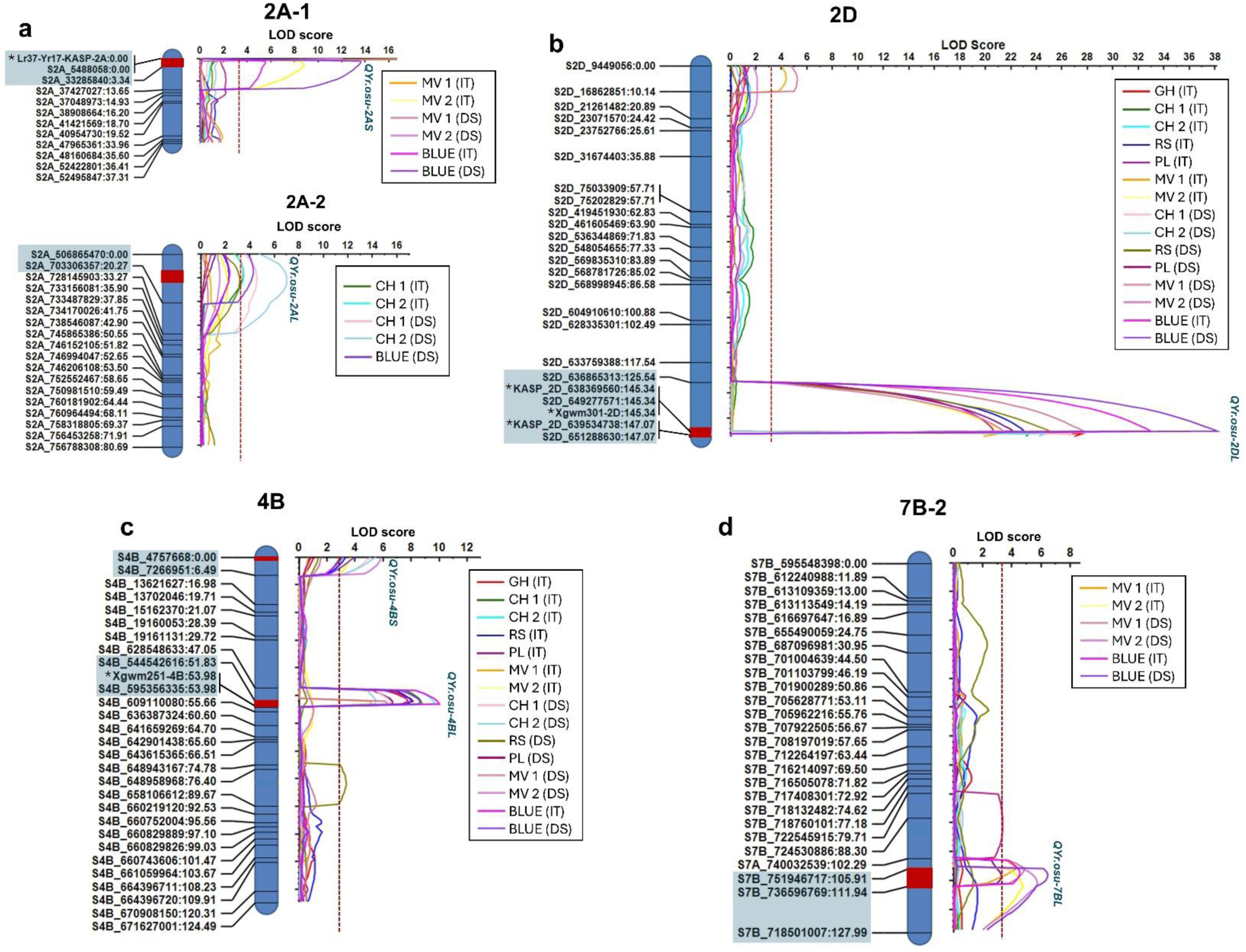
Genetic maps and quantitative trait loci (QTL) analysis identified six QTL detected in the doubled haploid population OK12D22004-016 × Baker’s Ann. a) The QTL *QYr.osu-2AS* and *QYr.osu-2AL* on chromosome 2A originated from OK12D22004-016. b) *QYr.osu-2DL* on chromosome 2D originated from Baker’s Ann. c) *QYr.osu-4BS* and *QYr.osu-4BL* on chromosome 4B originated from Baker’s Ann. d) *QYr.osu-7BL* on chromosome 7B originated from Baker’s Ann. Marker genetic positions are in centiMorgans (cM). Red rectangle on genetic map indicates the respective QTL. Highlighted markers in the genetic map correspond to flanking markers of the identified QTL. Red dashed line on logarithm of odds (LOD) curves indicates LOD threshold to determine QTL which was set at 3.3 at 1000 permutations with *P* = 0.05. IT = Infection type; DS = Disease severity (%); GH = Greenhouse (4–20°C); CH 1 = Chickasha, OK (evaluated on April 30), CH 2 = Chickasha, OK (evaluated on May 4); RS =Rossville, KS (evaluated on May 17); PL = Pullman, WA (evaluated on June 12); MV 1 = Mount Vernon (evaluated on April 10); MV 2 = Mount Vernon (evaluated on June 6); BLUE = Multi-environment best linear unbiased estimates. In genetic maps, markers with asterisk (*) are SSR or KASP markers previously linked with known *Yr* genes/QTL: *Lr37-Yr17-Sr38_GBG-KASP* linked to the 2N^v^S translocation (Liu et al., 2020), *Xgwm301* linked to *Yr54* (Basnet et al., 2014), *KASP_2D:638369560* linked to *QYr.hwwg-2D* (Mustahsan et al., 2023), *KASP_2D:639534738* linked to *QYr.hwwg-2D* (Mustahsan et al., 2023), and *Xgwm251* linked to *Yr62* (Lu et al., 2014).

**Table 2.**
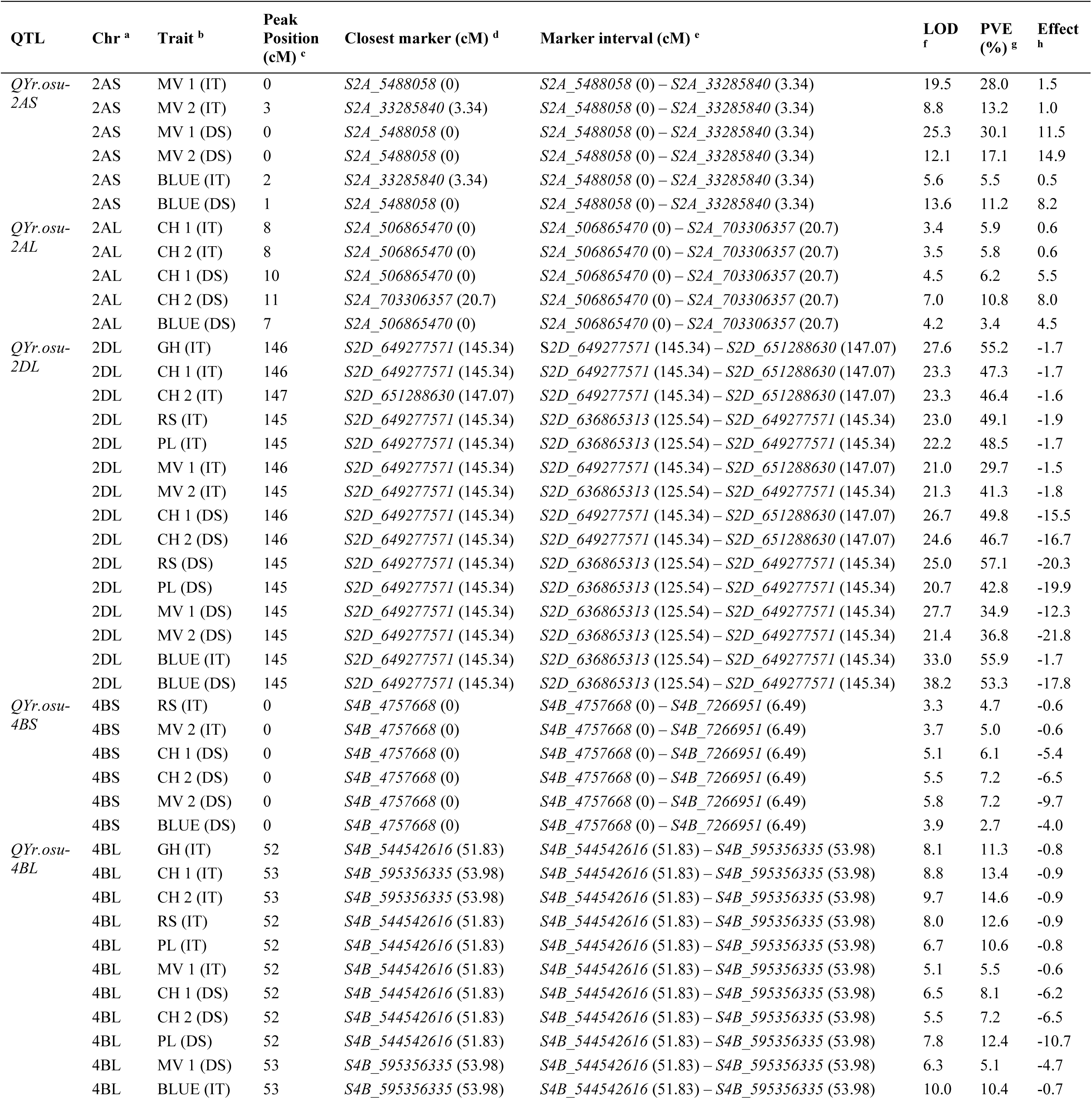
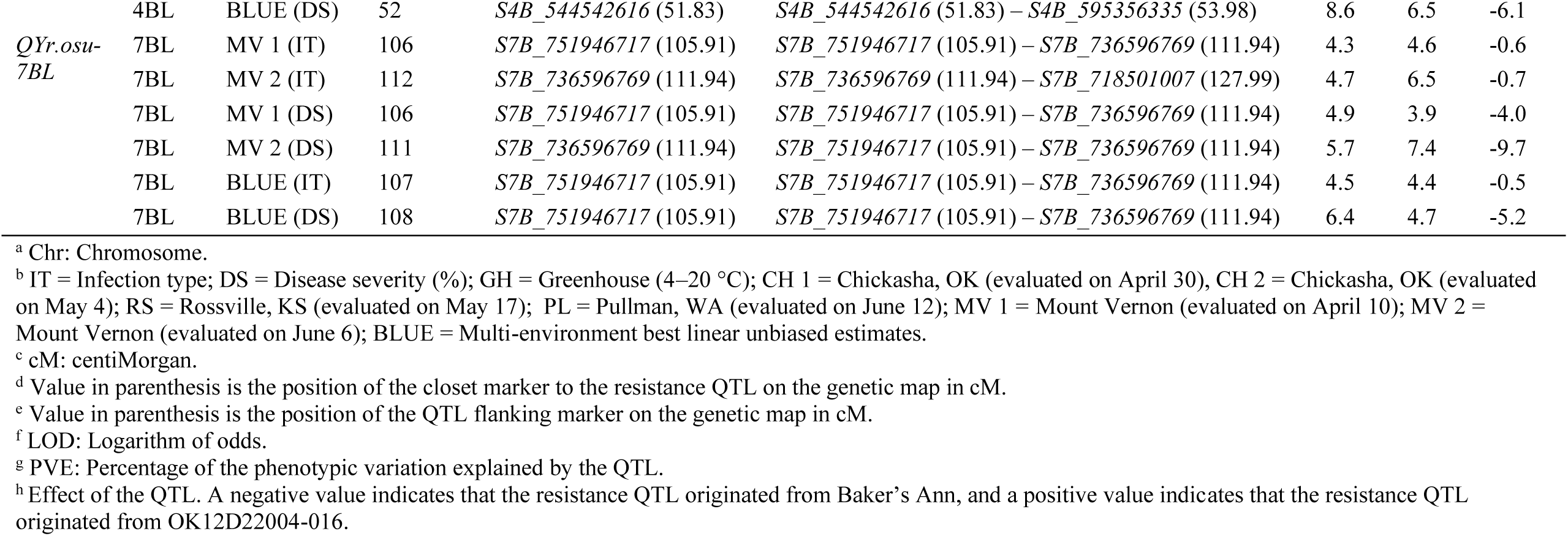
Summary of adult plant stripe rust resistance QTL detected in the doubled haploid population OK12D22004-016 × Baker’s Ann.

For OK12D22004-016, a major and a minor QTL were identified. The major QTL, *QYr.osu-2AS*, was mapped on the short arm of chromosome 2A within 5.4 to 33.2 Mb. This QTL was flanked by the markers *S2A_5488058* (0 cM) and *S2A_33285840* (3.34 cM) and was significant in MV 1 and MV 2 and BLUE with LOD scores of 5.6 to 25.3, explaining up to 30.1% of the phenotypic variation. The minor QTL, *QYr.osu-2AL*, was located on the long arm of chromosome 2A (506.8–703.3 Mb) and was flanked by the markers *S2A_506865470* and *S2A_703306357* (0 – 20.7 cM). This QTL was detected for IT and DS in CH 1, CH 2 and BLUE (DS), accounting for 3.4 to 10.8% of the phenotypic variation (Table 2; Figure 3).

### Development and validation of KASP markers

The two SNP markers, *S2D_649277571* and *S2D_651288630*, which are the closest to the major QTL in Baker’s Ann, *QYr.osu-2DL*, were targeted for the development of KASP markers. *KASP_ S2D_649277571* was successfully developed for the SNP *S2D_649277571*. However, a KASP marker could not be developed for the SNP *S2D_651288630* because its flanking sequence has high GC content. Therefore, *KASP_ S2D_650730433* was developed for another SNP marker, *S2D_650730433*, from the same bin (mapped at the same genetic position as *S2D_651288630*) (Table S8). The primer sequences for both KASP markers were provided in Table 3. The two KASP markers were validated on the parents and the 125 DH lines and were effective in differentiating the resistant and susceptible lines (Figure 4). Genotype calls for the two KASP markers agreed with their corresponding GBS-SNP marker data except for a single DH line, where *KASP_S2D_650730433* did not match the corresponding GBS-SNP call, likely due to a GBS genotyping error or a DNA contamination (Table S9).

**Figure 4.**
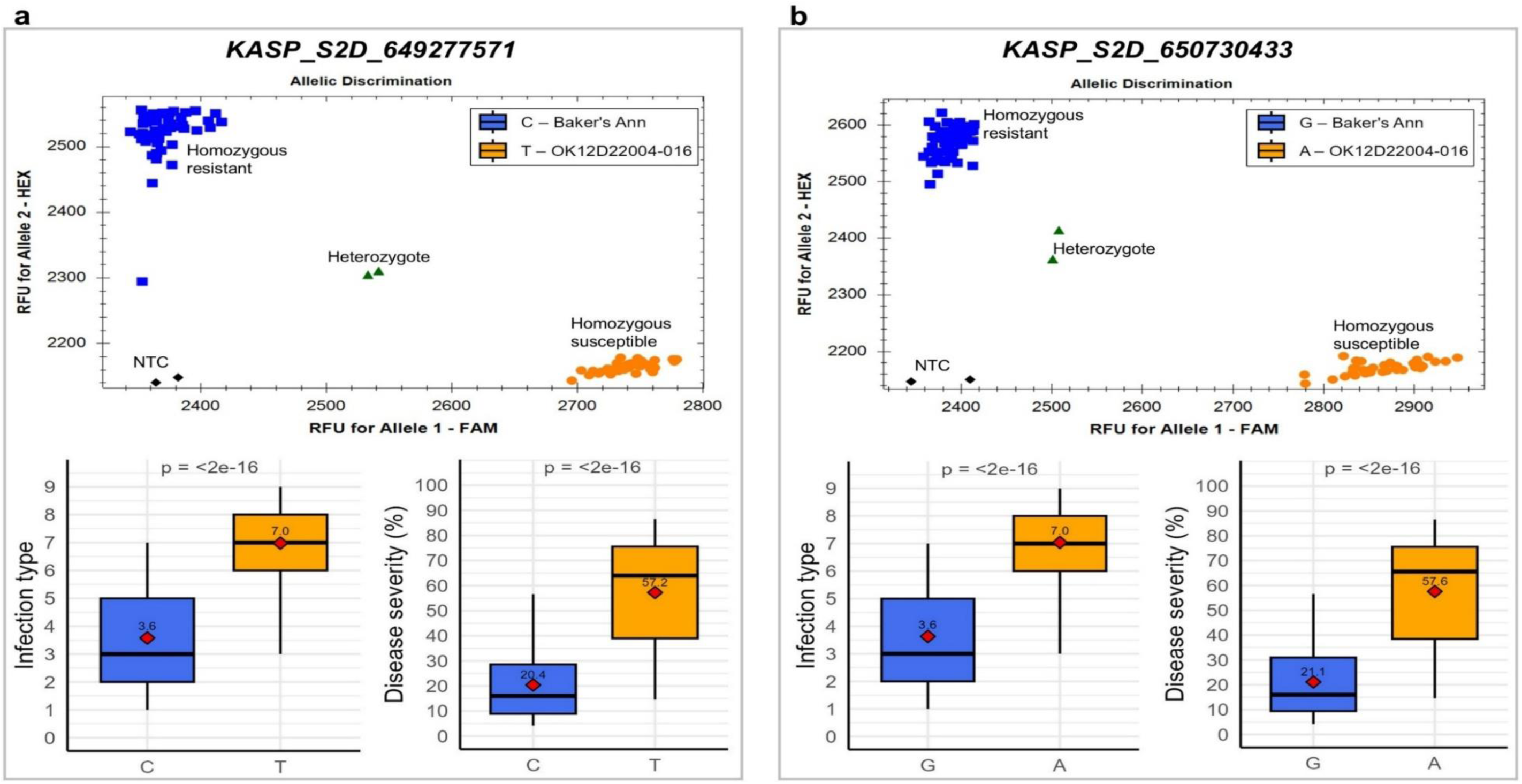
Genotyping scatterplots for the kompetitive allele-specific PCR (KASP) markers (a) *KASP_S2D_649277571* and (b) *KASP_S2D_650730433*, developed for the major QTL from Baker’s Ann, *QYr.osu-2DL* on chromosome 2D. Green triangles in scatterplots indicate artificial heterozygous controls included in the KASP assay. Below each scatterplot, there were box plots comparing stripe rust responses based on multi-environment BLUE of the doubled haploid lines carrying either the resistant or the susceptible allele. Horizontal lines within the box denote medians, and red diamonds represent the means.

**Table 3.**
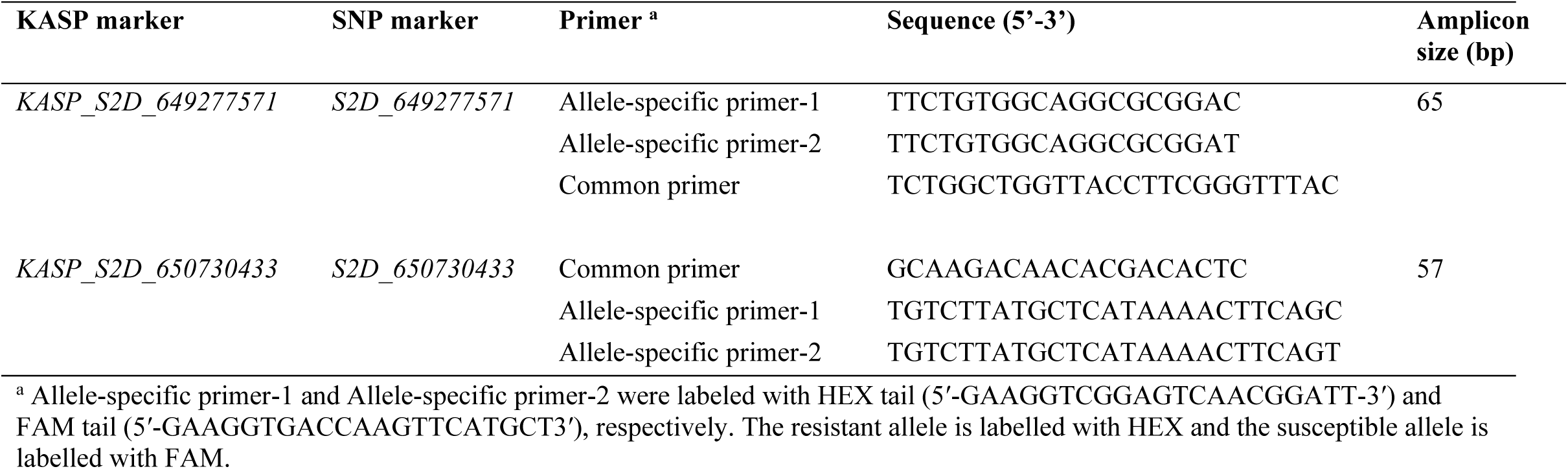
Kompetitive allele-specific PCR (KASP) markers developed from GBS SNP markers linked to the major QTL, QYr.osu-2DL, identified in Baker’s Ann.

### Effects of Baker’s Ann QTL combinations

The effects of individual QTL and their combinations were determined by classifying the DH lines into 19 groups. Based on BLUE data, lines with *QYr.osu-2DL* alone were moderately resistant (IT 5.1 and DS 32. 4%) (Figure 5). Based on Tukey’s HSD test, DH lines with either *QYr.osu-4BL*, *QYr.osu-4BL*, or *QYr.osu-7BL* had similar BLUE for IT and DS to that of DH lines with no QTL DH lines with *QYr.osu-2DL* and *QYr.osu-4BL* had significantly lower IT but not DS, compared to lines with only *QYr.osu-2DL*. Lines carrying all four QTL detected from Baker’s Ann (*QYr.osu-2DL*, *QYr.osu-4BS*, *QYr.osu-4BL*, and *QYr.osu-7BL*) showed the highest resistance level (Figure 5). ANOVA conducted on the four Baker’s Ann QTLs and their pairwise combinations revealed that while some QTL exhibited significant main effects, their interactions were not statistically significant, indicating that the QTL in Baker’s Ann are additive (Table S10).

**Figure 5.**
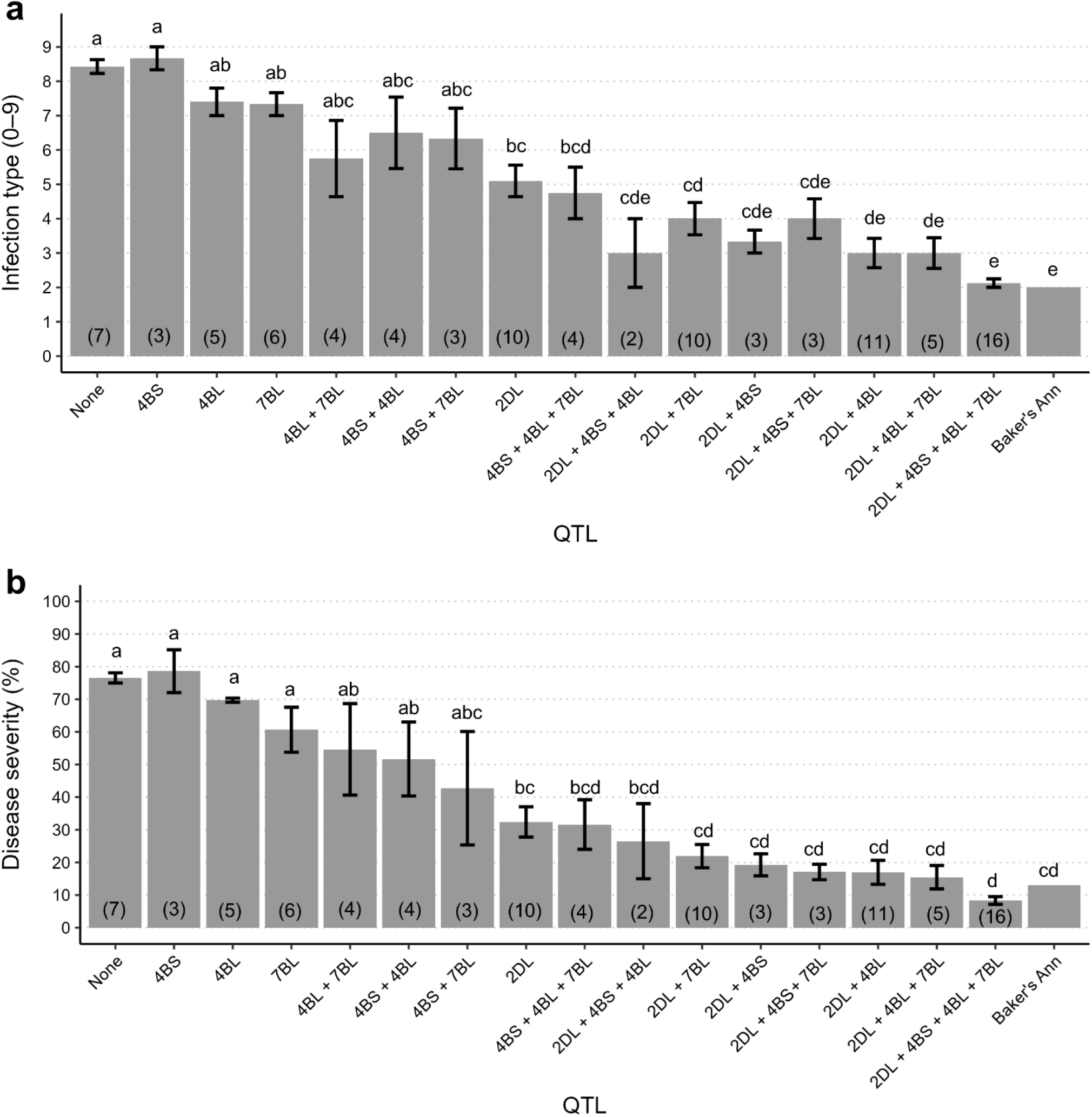
Effects of individual quantitative trait loci (QTL) and QTL combinations on stripe rust response in the OK12D22004-016 × Baker’s Ann doubled haploid population, based on multi-environment best linear unbiased estimates for (a) infection type and (b) disease severity. Numbers beneath each bar indicate the number of lines carrying that QTL combination. Bars labelled with different letters differ significantly based on Tukey’s HSD test.

## Discussion

Stripe rust is considered the most damaging wheat disease in the heart of the HWW growing region of the US Great Plains. This region experienced virulence changes in *Pst* races since the early 2000s that have compromised the resistance of most ASR genes and several APR genes (Hovmøller et al., 2011; Mu et al., 2020; Sørensen et al., 2014; Wan & Chen, 2014). Continuous breeding efforts to develop cultivars with durable stripe rust resistance largely depend on identifying and characterizing resistance genes. In this study, we identified the stripe rust resistance QTL in the HWW cultivar Baker’s Ann. The parents of the bi-parental cross, Baker’s Ann and OK12D22004-016 were evaluated in the Southern Regional Performance Nursery (SRPN) in 2017 and 2018, respectively (https://www.ars.usda.gov/plains-area/lincoln-ne/wheat-sorghum-and-forage-research/docs/hard-winter-wheat-regional-nursery-program/research/). Based on molecular markers data from the SRPN for seven characterized *Yr* genes (*Yr5*, *Yr15*, *Yr17*, *Yr18*, *Yr29*, *Yr36*, and *Yr46*), Baker’s Ann carries *Yr29*, whereas OK12D22004-016 carries *Yr5*, *Yr17*, and *Yr46*. The identification of *QYr.osu-2AS* in OK12D22004-016 confirmed the presence of the 2N^v^S translocation that carries *Yr17* in this breeding line. OK12D22004-016 is susceptible at the seedling stage to US *Pst* races, indicating that *Yr5*-linked marker (Naruoka et al., 2016) is not perfect and may provide false positives. A low frequency of false positives was also found to be associated with the *Yr5* marker, based on a panel of 459 HWW from the US Great Plains (Sharma et al. 2025). No QTL within the genomic region of *Yr46* was detected in Baker’s Ann based on this QTL analysis study. Sharma et al. (2025) found that *Yr46* is absent in contemporary US Great Plains HWW, so it is most likely that the marker linked to *Yr46* (Moore et al., 2015) provided a few false positives or it could be a genotyping error. In the present study, no QTL was detected within the genomic region of *Yr29*, indicating that the diagnostic power of the available *Yr29* marker (Brown-Guedira, G. & Fellers, J.P., unpublished) is limited. *Yr29* (same locus as the leaf rust resistance gene *Lr46*) was also not detected in genome-wide association studies (GWAS) for leaf rust and stripe rust resistance in US HWW genotypes, despite its presence based on molecular markers in 59% of the genotypes (Lakkakula et al., 2025; Sharma et al., 2025). This indicates high rate of false positive associated with the available *Lr46/Yr29* linked marker as this gene is not yet cloned.

*QYr.osu-2DL* was mapped to the telomere of chromosome 2DL and was consistently associated with stripe rust response across environments. In nine out of 15 environments, this QTL was flanked by the SNP markers *S2D_636865313* (636.7 Mb) and *S2D_649277571* (649.3 Mb), and in the remaining six environments between *S2D_649277571* (649.3 Mb) and *S2D_651288630* (651.3 Mb). For most environments, *S2D_649277571* was the closest to the QTL peak. Five designated *Yr* genes and three temporary named *Yr* genes have been identified on chromosome 2D. Among them, *Yr8*, *Yr37*, *YrCK*, *YrHRMSN-81*, and *YrH9020* are classified as ASR genes (Bariana et al., 2001; Chen et al., 2013; Li et al., 2012; Marais et al., 2005; Wang & Chen, 2017), thus unlikely to be associated with our identified APR QTL, *QYr.osu-2DL*. *Yr16* is an APR gene, identified in the cultivar “Capelle–Desprez” on the short arm of chromosome 2D (Agenbag et al., 2012), which is far from the position of *QYr.osu-2DL*. Another APR gene on chromosome 2DL, *Yr54*, was identified in the spring wheat line “Quaiu 3” and explained 49 – 54% of the phenotypic variation (Basnet et al., 2014). *Yr54* was flanked by the diversity arrays technology (DArT) markers *wpt-667162* and *wpt-667054*, which lies proximal to the SSR marker *Xgwm301* (648.9 Mb). *Xgwm301* was polymorphic between the parents of the bi-parental cross in this study and co-segregated with *QYr.osu-2DL* flanking marker *S2D_649277571* (Figure 3; Table S7). DH lines carrying only *QYr.osu-2DL* confers moderate resistance (IT 5.1 and DS 32.4%), consistent with that reported for *Yr54*. Therefore, *QYr.osu-2DL* identified in Baker’s Ann is likely *Yr54*. Further work is needed to identify the relationship between *QYr.osu-2DL* and *Yr54. Yr55* is another APR gene that was mapped within the genomic region of *QYr.osu-2DL* and flanked by the SSR markers *Xmag4089* (614.1 Mb) and *Xmag3385* (649.9 Mb) (McIntosh et al., 2014). However, both *Xmag4089* and *Xmag3385* were monomorphic for the parents of our bi-parental population, suggesting that *QYr.osu-2DL* is likely distinct from *Yr55.* Mustahsan et al. (2023) identified *QYr.hwwg-2D* in the HWW cultivar “Overland” between 650.6 and 650.7 Mb and explained 12.6 to 13.4% of phenotypic variation. *QYr.hwwg-2D* flanking markers, *KASP_2D:638369560* and *KASP_2D:639534738*, co-segregated with the flanking markers of *QYr.osu-2DL* in Baker’s Ann and *Yr54* linked marker *Xgwm301* (Figure 3; Table S7), suggesting that both QTL could be the same or linked. Exome capture for a set of 83 HWW genotypes, including Overland and Baker’s Ann identified 1,145 SNPs in the region of *QYr.osu-2DL* (636,865,313 bp to the end of chromosome 2DL telomere at 656,390,129 bp) (Table S11, https://doi.org/10.6084/m9.figshare.30024811.v4). Of these 1,145 SNPs, there were 164 polymorphic SNPs between Overland and Baker’s Ann. A close neighbor join and a high identity-by-state was identified between Overland and Baker’s Ann in this region, indicating that *QYr.osu-2DL* in Baker’s Ann and *QYr.hwwg-2D* in Overland might have the same origin. However, further investigation is needed to confirm their relationship. Another major APR QTL within the genomic region of *QYr.osu-2DL* was *QPst.jic-2D*, flanked by the SSR markers *Xgwm320* (646.9 Mb) and *Xgwm301* (648.9 Mb) (Jagger et al., 2011). Several loci associated with stripe rust at the adult plant stage and mapped close to *QYr.osu-2DL* were also identified in association mapping studies including, *Qyr.wpg*-*2D.2* (*IWA6851* [652.7 Mb] and *IWA5211* [652.9 Mb]), detected in US Pacific Northwest (PNW) winter wheat (Naruoka et al., 2015), *QYr.cim-2DL.2* tagged by marker *wpt-667054* (654.3 Mb) (Juliana et al., 2020), and an APR QTL associated with the SNP *1071080* (625.9 Mb) in wheat landraces (Tehseen et al., 2021).

Two QTL were detected on chromosome 4B in Baker’s Ann: a minor effect QTL, *QYr.osu-4BS* (4.7 – 7.3 Mb), located in the telomeric region of 4BS and a major QTL, *QYr.osu-4BL* (544.5 – 595.3 Mb), mapped on chromosome 4BL. The resistance genes, *Yr50*, *Yr62*, *Yr68*, *YrCH45-2*, and *YrR39* were mapped on chromosome 4B. However, *Yr50* and *YrCH45-2* provide ASR, with *Yr50* being introgressed from *Thinopyrum intermedium* into the Chinese cultivar “CH223” (Liu et al., 2013), and *YrCH45-2* mapped in the spring wheat cultivar “Chuanmai 45” on 4BL (Yang et al., 2016). Although *YrR39* confers APR, it was mapped between 65.5 and 243.2 Mb (Yin et al., 2018), which is far from the positions of *QYr.osu-4BS* and *QYr.osu-4BL. Yr62* and *Yr68*, were mapped within the physical position of *QYr.osu-4BL. QYr.osu-4BL* was identified in nearly every environment in this study. *Yr62* is a high-temperature adult-plant (HTAP) resistance gene, which was mapped in the Portuguese spring wheat “PI 192252” and flanked by the SSR markers, *Xgwm192* (508.1 Mb) and *Xgwm251* (567.7 Mb) (Lu et al., 2014). Integrating the flanking markers of *Yr62* in our mapping study showed that *Xgwm192* was monomorphic, whereas *Xgwm251* was polymorphic between the parents of the DH population and co-segregated with one of the flanking markers of *QYr.osu-4BL* (Figure 3; and Table S7). This indicates that *QYr.osu-4BL* is likely *Yr62* and further studies are needed to validate their relationship. Comparative mapping showed that the APR gene, *Yr68*, from the wheat line “AGG91587WHEA1”, which was mapped close to the SNP marker *IWA4640* (578.2 Mb) (McIntosh et al., 2016) was within the genomic region of *QYr.osu-4BL*. However, a developed KASP marker from *IWA4640* was found to be monomorphic between the parents of the DH population, suggesting that *QYr.osu-4BL* is likely different from *Yr68*. Previous QTL analysis studies identified APR QTL within the genomic region of *QYr.osu-4BS* including *QYr.hebau-4BS* in the spring wheat cultivars “Fuyu 3” at 5.0 Mb and “Thatcher” at 15.0 Mb (Gebrewahid et al., 2020; Tong et al., 2024; Zhang et al., 2019). A GWAS study by Sharma et al. (2025) identified a significant association between the SNP marker *S4B_571886653* (571.9 Mb) and stripe rust response in US Great Plains HWW, which was within the physical position of *QYr.osu-4BL*.

Another minor effect QTL from Baker’s Ann, *QYr.osu-7BL*, was detected for stripe rust response in Mount Vernon and BLUE. It was mapped near the distal end of chromosome 7BL (718.5 – 751.9 Mb), a region rich in *Yr* genes, most of which confer ASR. Four HTAP genes, *Yr39*, *Yr52*, *Yr59*, and *Yr79* were mapped on chromosome 7BL in the US spring wheat cultivar “Alpowa” (Lin & Chen, 2007), the Indian line “PI 183527” (Ren et al., 2012), the Iraqi line “PI 178759” (Zhou et al., 2014) and the Pakistani landrace “PI 182103” (Feng et al., 2018), respectively. Based on physical positions, *Yr39* (609.6 Mb) and *Yr79* (227.8 – 247.1 Mb) lie outside the physical interval of *QYr.osu-7BL*, whereas *Yr52* closely linked to *Xbarc182* (744.2 Mb), and *Yr59* linked to *Xbarc32* (734.1 Mb) were within the genomic region of *QYr.osu-7BL*. The relationship between *QYr.osu-7BL* and these two APR genes (*Yr52* and *Yr59*) remains unclear and will require further investigation.

Two QTL were identified in OK12D22004-016, a major QTL *QYr.osu-2AS* (5.5 – 33.3 Mb) on chromosome 2AS, and a minor effect QTL *QYr.osu-2AL* (506.7 – 703.3 Mb) on chromosome 2AL. *QYr.osu-2AS* is associated with the 2N^v^S translocation that carries *Yr17*. *Lr37-Yr17-Sr38_GBG-KASP* (Liu et al., 2020) was polymorphic in the current mapping population and co-segregated with *QYr.osu-2AS* flanking marker, *S2A_5488058* (Figure 3; and Table S7). *Yr17* along with 2N^v^S segment was translocated from *Aegilops ventricosa* into the hexaploid wheat line “VPM 1” (Bariana & McIntosh, 1993), and this 33 Mb segment does not recombine with hexaploid wheat genome (Gao et al., 2021). The source of the 2N^v^S introgression in OK12D22004-016 is most likely the HWW cultivar “Overley”, which appears in its pedigree. Mustahsan et al. (2023) reported *QYr.hwwg-2A* in Overley within the 2N^v^S region. Interestingly, *QYr.osu-2AS* was detected only in Mount Vernon and BLUE data. Since most US *Pst* races, including the most common race in the US PSTv-37, are virulent to *Yr17* (Wang et al., 2022), this QTL is unlikely *Yr17*. A HTAP resistance gene, *YrM1225*, has been mapped on the 2N^v^S/2AS segment, which co-segregated with marker *Lr37-Yr17-Sr38_GBG-KASP* (Li et al., 2023). Thus, *YrM1225* or other genes on the 2N^v^S translocation likely account for the resistance associated with *QYr.osu-2AS*. The minor effect QTL from OK12D22004-016, *QYr.osu-2AL* (506.8 – 703.3 Mb), was identified for stripe rust responses in Chickasha and with the BLUE. The APR genes *Yr86* (710.3 – 712.6 Mb) (Cao et al., 2024; Zhu et al., 2023) and *Yrxy2* linked with *Xbarc5* at 678.1 Mb (Zhou et al., 2011) were within the genomic region of *QYr.osu-2AL.* Previously identified QTL on chromosome 2AL that were mapped close to *QYr.osu-2AL* include *QYrWS.wgp-2AL* (611.6 – 684.7 Mb) (Upadhaya et al., 2025) and *QYrPI197734.wgp-2A* (559.7 – 713.4 Mb) (Liu et al., 2020). However, *QYr.osu-2AL* was detected in Chickasha, whereas *QYrWS.wgp-2AL* and *QYrPI197734.wgp-2A* was detected in PNW environments. Consequently, it is difficult to determine the relationship of *QYr.osu-2AL* with previously reported *Yr* genes or QTL in this region.

In conclusion, Baker’s Ann has a strong APR to stripe rust along with superior agronomic and end-use quality traits. Four QTL located on chromosomes 2DL, 4BS, 4BL, and 7BL explained its high APR to stripe rust. Two KASP markers, *KASP_ S2D_649277571* and *KASP_ S2D_650730433* linked to the major QTL in Baker’s Ann, *QYr.osu-2DL*, were developed to help the transfer of this resistance source into future breeding lines. *QYr.osu-2DL* was found in proximity to the APR gene *Yr54* and further investigation is needed to validate their relationship. A second major QTL in Baker’s Ann was *QYr.osu-4BL,* which is likely *Yr62.* Although OK12D22004-016 is susceptible in the US Great Plains, it was found to possibly carry an APR gene on the 2N^v^S translocation, which was effective in the PNW and can enhance stripe rust resistance when pyramided with other resistance genes.

## Data availability statement

All data generated or analyzed during this study are included in this published article and its supplementary information files submitted with this manuscript. The raw sequencing data can be accessed from the NCBI Short-Read Archive BioProject PRJNA1312051 (ncbi.nlm.nih.gov/sra/PRJNA1312051). Supplemental tables (Table S1-S10) are available at https://doi.org/10.6084/m9.figshare.30047773.v1. Table S11. Exome capture SNP data for 83 hard winter wheat accessions in the region of *QYr.osu-2DL* (Supplemental Table S11) is available at https://doi.org/10.6084/m9.figshare.30024811.v4.

## Acknowledgments

This project was funded by the US Department of Agriculture National Institute of Food and Agriculture Grant # 2024-67013-42587. The mention of trade names or commercial products in this publication is solely to provide specific information and does not imply recommendation or endorsement by the US Department of Agriculture. The USDA is an equal opportunity provider and employer.

## Supplemental Material

### Supplemental Tables

**Supplemental Table S1.** GBS-SNP marker data for the doubled haploid population OK12D22004-016 × Baker’s Ann, including 13,090 SNP markers prior to filtering.

**Supplemental Table S2.** Genotypic data for the doubled haploid population OK12D22004-016 × Baker’s Ann, including 7,268 SNP markers and five KASP/SSR markers linked to known *Yr* genes/QTL.

**Supplemental Table S3.** Primer sequences of SSR and KASP markers linked to stripe rust resistance genes/QTL in proximity to the identified major QTL in our study.

**Supplemental Table S4.** Stripe rust responses at the adult plant stage of 125 doubled haploid lines of the cross OK12D22004-016 × Baker’s Ann.

**Supplemental Table S5.** Analysis of variance for stripe rust infection type and disease severity in the OK12D22004-016 × Baker’s Ann doubled haploid population evaluated across environments.

**Supplemental Table S6.** Summary of the linkage mapping using 661 molecular markers for the bi-parental cross OK12D22004-016 × Baker’s Ann.

**Supplemental Table S7.** Molecular markers (excluding redundant markers) mapped to wheat chromosomes in the doubled haploid population OK12D22004-016 x Baker’s Ann.

**Supplemental Table S8**. Linkage map of chromosome 2D constructed using 265 markers for the doubled haploid population OK12D22004-016 x Baker’s Ann.

**Supplemental Table S9.** Comparison of genotyping calls of KASP markers and their corresponding GBS SNPs flanking *QYr.osu-2DL* from Baker’s Ann.

**Supplemental Table S10.** Analysis of variance for infection-type and disease-severity, conducted on the four Baker’s Ann QTL and their pairwise interactions.

**Supplemental Table S11**. Exome capture SNP data for 83 hard winter wheat accessions in the region of *QYr.osu-2DL*.

### Supplemental Figure

**Supplemental Figure S1.** Correlations between stripe rust responses in different environments at the adult plant stage for the doubled haploid population OK12D22004-016 × Baker’s Ann.

## Abbreviations

ABC: ATP-binding cassette
APR: adult plant resistance
ASR: all-stage resistance
BLUE: best linear unbiased estimates
cM: centiMorgan
CTAB: cetyltrimethylammonium bromide
DArT: diversity arrays technology
DH: doubled haploid
DS: disease severity
GBS: genotyping-by-sequencing
MAS: marker-assisted selection
GWAS: genome-wide association studies
HTAP: high-temperature adult-plant
HWW: hard winter wheat
ICIM-ADD: inclusive composite interval mapping for additive quantitative trait loci
IT: infection type
KASP: kompetitive allele-specific PCR
LOD: logarithm of odds
NBS-LRR: nucleotide-binding site leucine-rich repeat
OSU: Oklahoma State University
PNW: Pacific Northwest
*Pst*: *Puccinia striiformis* f. sp. *tritici*
QTL: quantitative trait loci
SNPs: single-nucleotide polymorphisms
SRPN: Southern Regional Performance Nursery
SSR: simple sequence repeat
US: United States
*Yr*: stripe rust resistance gene

