## Supplemental Figure S1 for "Identification of stripe rust adult plant resistance genes in the hard winter wheat cultivar Baker’s Ann"

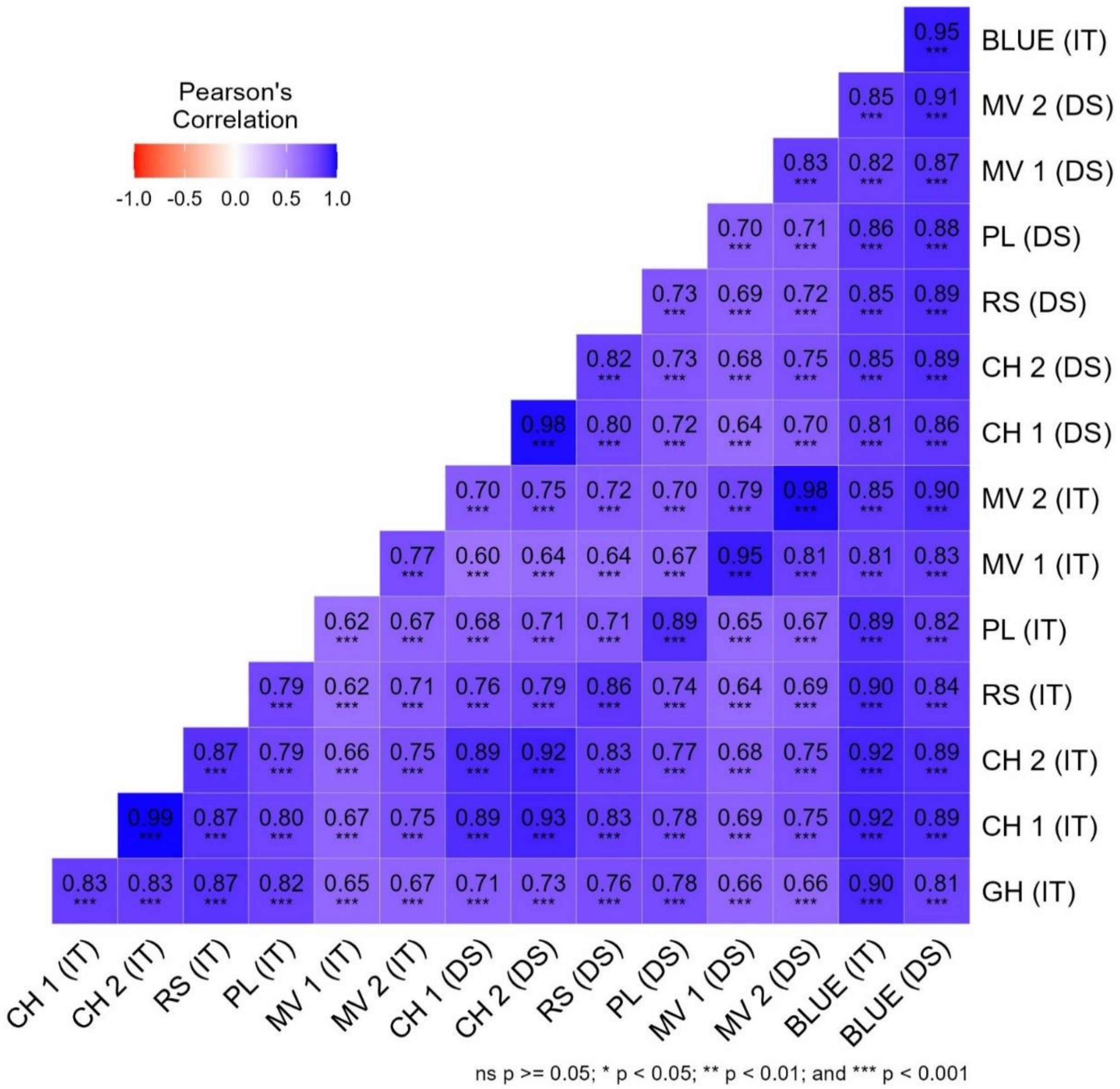


**Supplemental Figure S1**. Correlations between stripe rust responses in different environments at the adult plant stage for the doubled haploid population OK12D22004-016 x Baker's Ann. IT = Infection type; DS = Disease severity (%); GH = Greenhouse (4–20°C); CH 1 = Chickasha, OK (evaluated on April 30), CH 2 = Chickasha, OK (evaluated on May 4); RS =Rossville, KS (evaluated on May 17); PL = Pullman, WA (evaluated on June 12); MV 1 = Mount Vernon (evaluated on April 10); MV 2 = Mount Vernon (evaluated on June 6); BLUE = Multi-environment best linear unbiased estimates.
